# Hierarchical effects of pro-inflammatory cytokines on the post-influenza susceptibility to pneumococcal coinfection

**DOI:** 10.1101/060293

**Authors:** Stefanie Duvigneau, Niharika Sharma-Chawla, Alessandro Boianelli, Sabine Stegemann-Koniszewski, Van Kinh Nguyen, Dunja Bruder, Esteban A. Hernandez-Vargas

## Abstract

In the course of influenza A virus (IAV) infections, a secondary bacterial infection frequently leads to serious respiratory conditions provoking high hospitalization and death tolls. Although abundant pro-inflammatory responses have been reported as key contributing factors for these severe dual infections, the relative contribution of cytokines remain largely unclear.

In the current study, mathematical modelling on *in vivo* experimental data highlight IFN-*γ* as a decisive candidate responsible for impaired bacterial clearance, thereby promoting bacterial growth and systemic dissemination during acute IAV infection. Moreover, we found a time-dependent detrimental role of IL-6 in curtailing bacterial outgrowth which was however not as distinct as for IFN-*γ*. Importantly, our results furthermore challenge current beliefs that the TNF-*α* response or the increased availability of nutrients modulated by IAV infection have a central role to the bacterial outgrowth. Ultimately, our findings contribute to a detailed understanding of the mechanisms underlying impaired bacterial clearance following influenza infection.

## Introduction

Retrospective studies performed on victims of the devastating 1918/1919 influenza A virus (IAV) pandemic and also the recent H1N1 IAV pandemic revealed a high incidence of coinfections with unrelated bacterial pathogens [1–5]. In fact, 71% of the alarmingly high death toll during the 1918/1919 outbreak was attributed to coinfection with *Streptococcus pneumoniae* (*S. pneumoniae*) [5]. This copathogen is a human adapted gram-positive colonizer of the nasopharynx in asymptomatic children and individuals over 65 years of age but at the same time remains to be the most common cause of community-acquired pneumonia [6]. Animal and human studies have shown that preceding IAV infection enhances all aspects of *S. pneumoniae* pathogenesis from nasopharyngeal colonization to invasive pneumococcal disease [7], leading to the strong predisposition to lethal secondary pneumococcal infection in IAV infected patients.

Several mechanisms have been implicated in the viral-bacterial synergism which altogether demonstrate a multifactorial and complex nature of copathogenesis. One central dogma is the disruption of the protective alveolar epithelial cell barrier due to the cytolytic mode of viral replication which in turn exposes otherwise cryptic bacterial adherence factors on the basal membrane and thereby promotes invasive pneumococcal disease. Along these lines, a recent study showed that an increase in nutrient availability due to the virus-mediated accumulation of sialylated mucins and enhanced desialylation of host glycoconjugates in the upper respiratory tract was a major factor contributing to bacterial outgrowth [8].

Additionally identified and more debatable mechanisms are the IAV-mediated immune modulations such as immune cell dysfunction and apoptosis causing an aberrant production of inflammatory mediators in the case of a secondary bacterial encounter. Experimental reports indicate dampened innate inflammatory responses to the bacteria in IAV pre-infected hosts due to an enhanced activation threshold of lung innate immune cells that renders them hypo-responsive [9].

In contrast, a number of studies describe a massive and overshooting inflammatory cell influx due to the hyper-production of pro-inflammatory cytokines such as Type I interferons (IFN-I), Interferon-*γ* (IFN-*γ*), Interleukin-6 (IL-6) and Tumor Necrosis Factor-*α* (TNF-*α*) during secondary bacterial infection. These are often linked to pulmonary edema due to irreparable damage to the alveoli and immunopathology leading to mortality during coinfections [7,10–13]. Taken together, these studies strongly reflect an exacerbated cytokine and chemokine production to significantly contribute to the detrimental changes in the lung microenvironment that favor secondary bacterial infections. However it is still largely unknown what the relative contribution of these cytokines to bacterial outgrowth, morbidity and mortality in secondary infection is and whether they work alone or in synergism as ‘friend or foe’ to the coinfected host.

Dissecting the detailed contributions of the identified players in enhancing susceptibility to severe secondary bacterial disease following IAV as well as their respective interactions is crucial to develop prophylactic and therapeutic strategies. In the past, mathematical modelling has made valuable contributions to our understanding of IAV infection, focusing either on IAV replication [14–19] and/or host immune responses to IAV [20–29]. Regarding the interactions between IAV and *S. pneumoniae*, the only modeling approach proposed so far has been through the pioneering work of Smith *et al.* [30]. However, to the best of the authors knowledge, untangling the contributions of the different mechanisms by which changes in the immune response affect bacterial clearance in a temporal manner has not been attempted until now. Therefore, by combining the results of tailored *in vivo* experiments and mathematical modelling approaches, we aimed at clarifying the relative contributions of different underlying mechanisms of the the IAV-S. *pneumoniae* synergism.

## Results

**Study Design.** The dynamics of IAV and *S. pneumoniae* coinfection were investigated by establishing a murine model displaying disease upon subsequent infection with sub-lethal infection dosages of both copathogens. Secondary infection with 1×10^6^ CFU of *Streptococcus pneumoniae* strain TIGR4 (T4) was performed on day 7 after IAV infection based on previous experimental observations that indicated peak susceptibility to pneumococcal disease at this time point during acute IAV infection [3, 31]. Bacterial burden, viral titers, cytokine concentrations and alveolar macrophages (AM) counts were determined in the respiratory tract for three experimental groups: coinfected (IAV+T4), single IAV and single T4 infected animals. A schematic representation of the experiments is provided in Figure 1.

**Figure 1.**
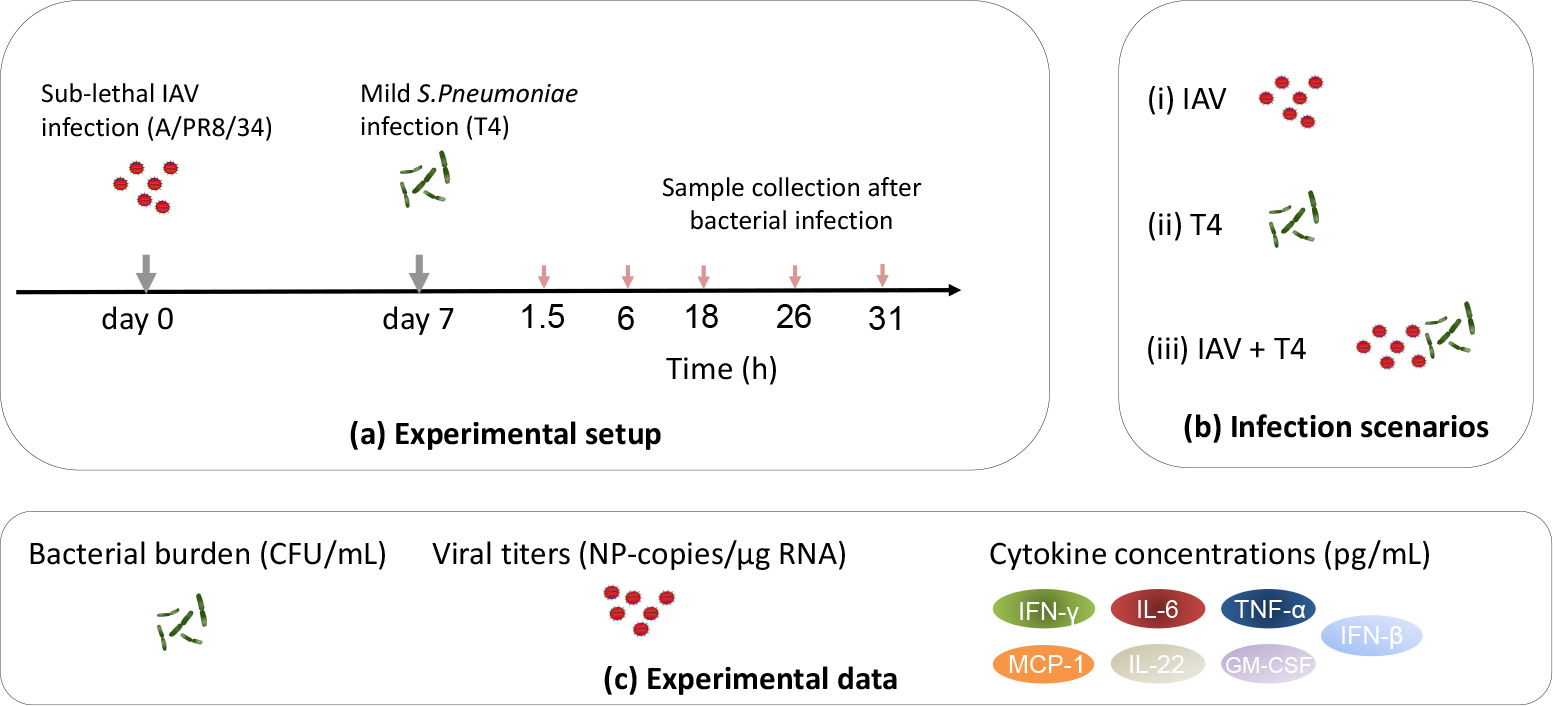
Experimental scheme. (**a**) C57BL/6J wildtype mice were intranasally infected with a sub-lethal dose of IAV (A/PR8/34) followed by bacterial infection with the *S. pneumoniae* strain T4 on day 7. BAL, post-lavage lung and blood were collected at indicated time points post secondary bacterial infection (hours post infection (hpi)). (**b**) The infection groups were single viral infection (IAV), single bacterial infection (T4) and coinfection (IAV+T4). (**c**) The bacterial burden, viral titers and cytokine concentrations were determined as the experimental readouts.

The complexity and at times redundancy of immune responses to infections often render to arduous and expensive experimental settings when attempting to identify the key components and their temporal contributions during coinfections. Thus, merging mathematical modelling with the relevant in *vivo* data is a promising tool to tailor future experiments [23,30,32,33]. In order to dissect the dynamics observed in our experimental results, mathematical modelling was employed not as a quantitative recapitulation of experimental data but as a tool to accept or reject hypotheses on the basis of various mathematical models as “thought experiments” using the Corrected Akaike Information Criterion (AICc) for the model selection process.

**Bacterial growth kinetics during IAV-pneumococcal coinfection.** Mouse coinfection experiments revealed that the bacterial load in the post-lavage lung tissue and bronchoalveolar lavage (BAL) were comparable between the IAV+T4 and single T4 infected groups until 6 hpi (Figure 2a and b). At 18 hpi, a significantly higher bacterial load was observed in both the lung tissue and BAL of the coinfected compared to the single T4 infected group. The bacterial load further increased until 31 hpi in the coinfected group. At the later time points, the high grade pneumonia, *i.e* the high bacterial numbers in the respiratory tract, was homogeneously accompanied by the systemic spread via the blood (bacteremia) in all the coinfected animals (Figure 2c). Significantly lower grade bacteremia was observed in the single T4 infected mice starting 18 hpi (Figure 2c).

Taken together, assessing the kinetics of bacterial growth and clearance in the respiratory tract and blood following IAV/*S. pneumoniae* coinfection revealed a “turning-point” between 6 and 18 hpi. At 18 hpi bacterial outgrowth became clearly evident in the coinfected group and was in strong contrast to the onset of bacterial clearance in the T4 only infected group (also see CFU data for individual mice in Supplementary Figure A.1).

**Figure 2.**
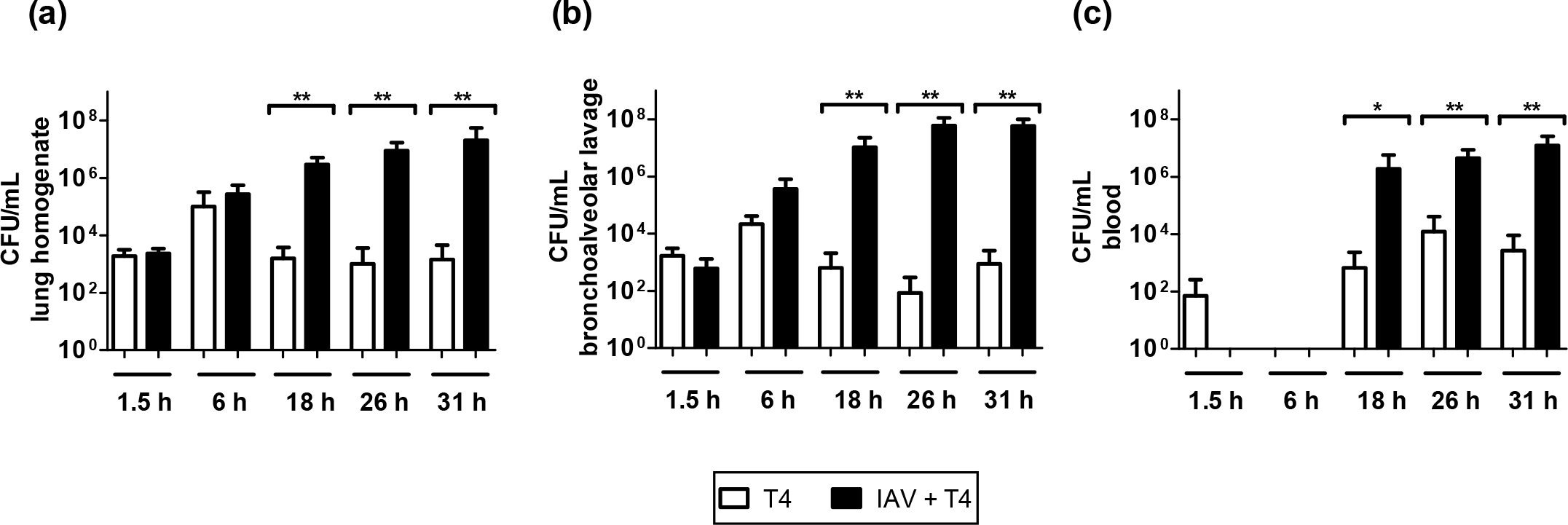
Organ-wide bacterial burden in the single and coinfected animals. Bacterial titers in single T4 infected and IAV+T4 coinfected mice were determined in (a) post-lavage lung, (b) BAL and (c) blood. All experiments were performed in groups of 4-7 WT C57BL/6J mice, raw data can be found in the Supplementary Figure A.1. Statistical analysis was performed using the two-tailed Mann-Whitney t test. Asterisks indicate significant differences between single and coinfected mice: *, p<0.05; **, p<0.01.

**Mathematical modeling revealed AM dynamics in the lungs of the coinfected animals insufficient to determine bacterial outgrowth.** Previous results in [34] revealed that 90% of resident AM are depleted in the first week after influenza infection forming a favourable niche for a secondary pneumococcal infection. To evaluate this finding in our experimental setting and mathematical modeling approach, we determined the absolute numbers of AM in the lung tissue early after secondary *S. pneumoniae* infection. Experimental results in Figure 3 show a significant decrease in AM numbers in the IAV+T4 group compared to the single T4 group at 18 hpi, coinciding with the established bacterial outgrowth in these mice at this time point.

**Figure 3.**
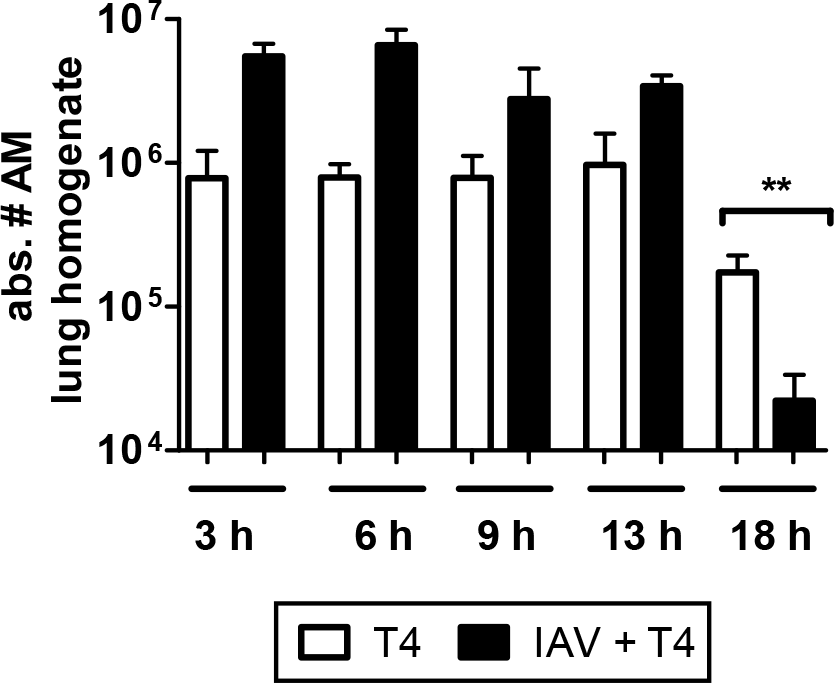
Absolute numbers of alveolar macrophages (AM) in post-lavage lungs of single and coinfected animals. Statistical analysis was performed using the two-tailed Mann-Whitney t test. Asterisks indicate significant differences between single and coinfected mice: **, p<0.01. Experiments were performed in groups of 3-4 WT C57BL/6J mice.

Thus, we evaluated the hypothesis of AM dynamics as a determinant to represent the bacterial kinetics in the single *S. pneumoniae* infected or in the coinfected group. To this end, by using AM counts dynamics for the different groups, we proposed several mathematical models to represent the dynamics observed in the single *S. pneumoniae* infected (Supplementary D) and coinfected animals (Supplementary E).

A model selection process by AICc revealed that assuming bacterial clearance with a constant number of functional AM (model D2 or D3 at Supplementary D) provided the best fitting to the data in the single T4 group. Surprisingly, for both the single T4 group and the IAV+T4 group, considering dynamic AM numbers rendered the worst fitting among the different models, thereby rejecting the hypothesis that bacterial clearance is mainly driven by AM dynamics. However, it is possible that only a fraction of functional AMs is required to clear the bacterial infection. These results were supported by considering the experimental data variability in mathematical models with bootstrapping procedures (see Supplementary F).

**Coinfection leads to a significant increase of IFN-*γ*, IL-6 and TNF-*α* airway concentrations.** In contrast to previous animal studies that often focused on a single time point post secondary infection [10, 11], we assessed the early kinetics of pro-inflammatory cytokines in the respiratory tract. For this purpose, we determined the IFN-*γ*, TNF-*α*, IL-6, Monocyte chemoattractant protein-1 (MCP-1), Interferon-*β* (IFN-*β*), Interleukin 22 (IL-22) and Granulocyte-macrophage colony-stimulating factor (GM-CSF) protein concentrations in the BAL fluid of single and coinfected mice. A time-dependent significant increase in the protein concentrations of IFN-*γ*, TNF-*α*, and IL-6 was observed in the coinfected animals when compared to the single T4 infected animals (Figure 4).

**Figure 4.**
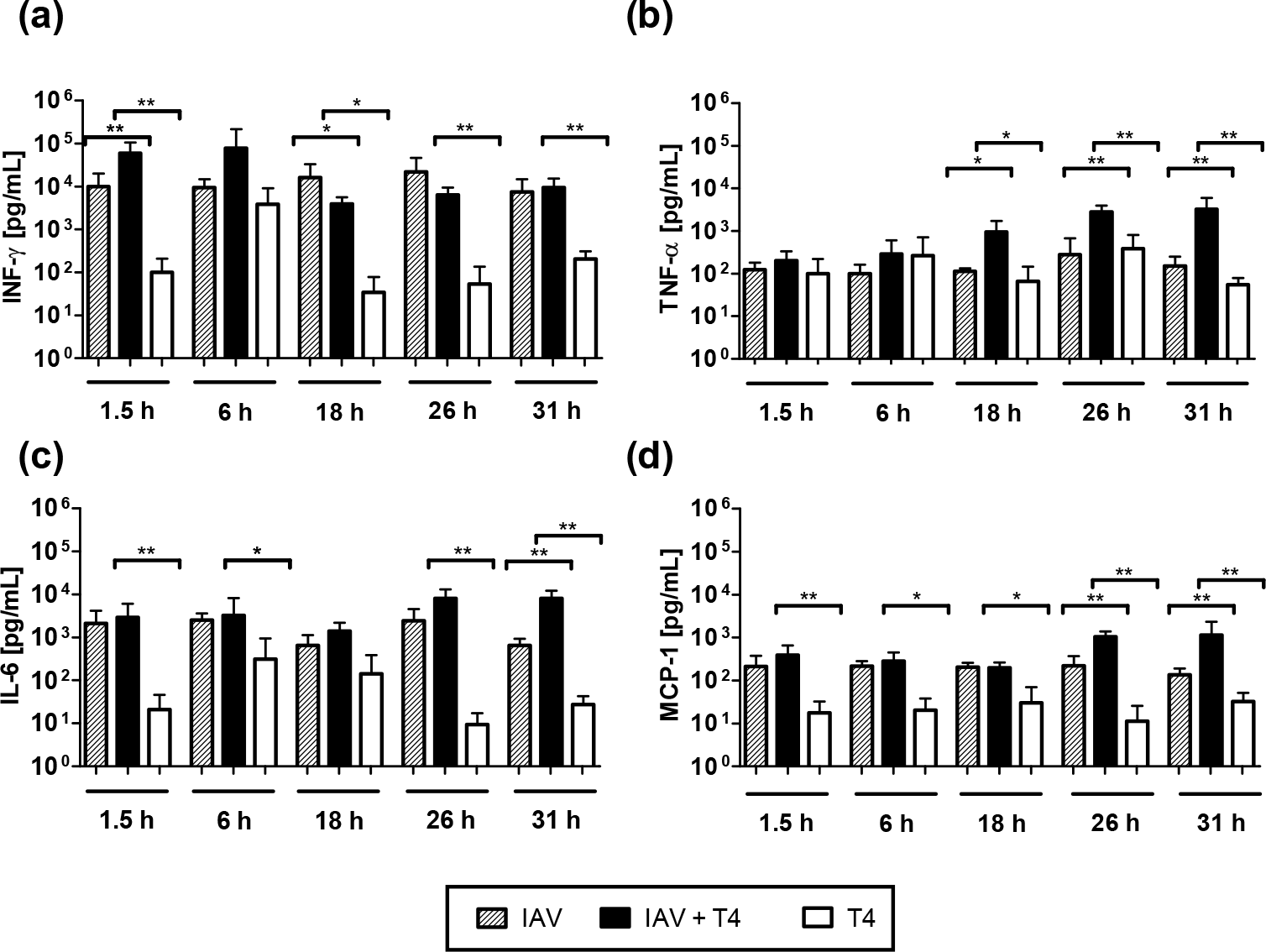
Pro-inflammatory cytokine profiles of the BAL of coinfected and single T4 infected mice. Protein concentrations of (a) IFN-*γ*, (b) TNF-*α*, (c) IL-6 and (d) MCP-1 were determined in the BAL fluid at the indicated time points after secondary T4 infection on day 7 post IAV or single T4 infection on day 7 post PBS treatment. Raw data can be found in the Supplementary Figure A.2. Asterisks indicate significant differences between single and coinfected mice: *, p<0.05; **, p<0.01.

Bacterial infection alone led to a transient increase in IFN-*γ* at 6 hpi and IL-6 at 6 hpi and 18 hpi, which hardly reached the elevated IFN-*γ* and IL-6 levels detected in the IAV-infected group at all time points analyzed (Figure 4). For IFN-*γ*, coinfection led a further increase early at 1.5 hpi and 6 hpi compared to the single IAV infection whereas the levels remained constant compared to the underlying IAV infection for the later time points and significant increase was only observed when compared to the single T4 infection. Overshooting IL-6 responses in the coinfected mice were detected at 26 hpi and 31 hpi compared to the single T4 infection. At the same time, TNF-*α* levels hardly changed between mice infected with IAV alone and T4 alone but were increasingly and significantly elevated in coinfected mice from 18 hpi on. This late surplus in IL-6 and TNF-*α* compared to both the single IAV and T4 infected groups indicated a steady induction of these two pro-inflammatory cytokines in the dual-infected animals (Figure 4b and c). Of note, the chemokine MCP-1 was also significantly increased in the IAV+T4 group compared to the single T4 infected group and marginally increased to the IAV only group at 26 hpi and 31 hpi (Figure 4d). The protein concentrations of the other inflammatory mediators did not show significant changes between the groups at all time points (see Supplementary Figure A.3).

**Decisive and robust role of IFN-*γ* in bacterial outgrowth.** To dissect the temporal contribution of the measured pro-inflammatory cytokines in preventing bacterial clearance, mathematical models fitted from single *S. pneumoniae* infection were challenged in order to assess the effects of pro-inflammatory responses on bacterial lung titers. Hence, we adopted the best model for single *S. pneumoniae* infection (model D2 from the Supplementary D) and challenged the mathematical term representing bacterial clearance (*c_b_B*) with different functions (*c_b_f_x_B*) to evaluate which of the pro-inflammatory cytokines or their combinations rendered the best fitting to the bacterial burden detected in the coinfected mice. A brief list of mathematical models tested is presented in Table 1 and a complete version with estimated parameters and parameter uncertainty analysis is shown in Supplementary E and F, respectively.

Considering the AICc scores, criterion of small differences (less than 2 units) were not significant (see Materials and Methods), the best group of models were M3 and M7 (Table 1). The common component of these two models are the IFN-*γ* kinetics and, remarkably, a mechanism solemnly based on the IFN-*γ* response (M3) provided a better fitting than both TNF-*α* (M4) and IL-6 (M5) even though we assumed a more conservative AICc criterion (*e.g* ≤ 10). Additionally, the models M6, M7, M8 and M9, that also had IFN-*γ* dynamics involved in the impairment function *f_x_*, scored closely to M3 (see Supplementary E). In agreement with previous works [11], our model selection process rendered IFN-*γ* as the crucial and sufficient modulator in the impairment of bacterial clearance.

**Table 1.**
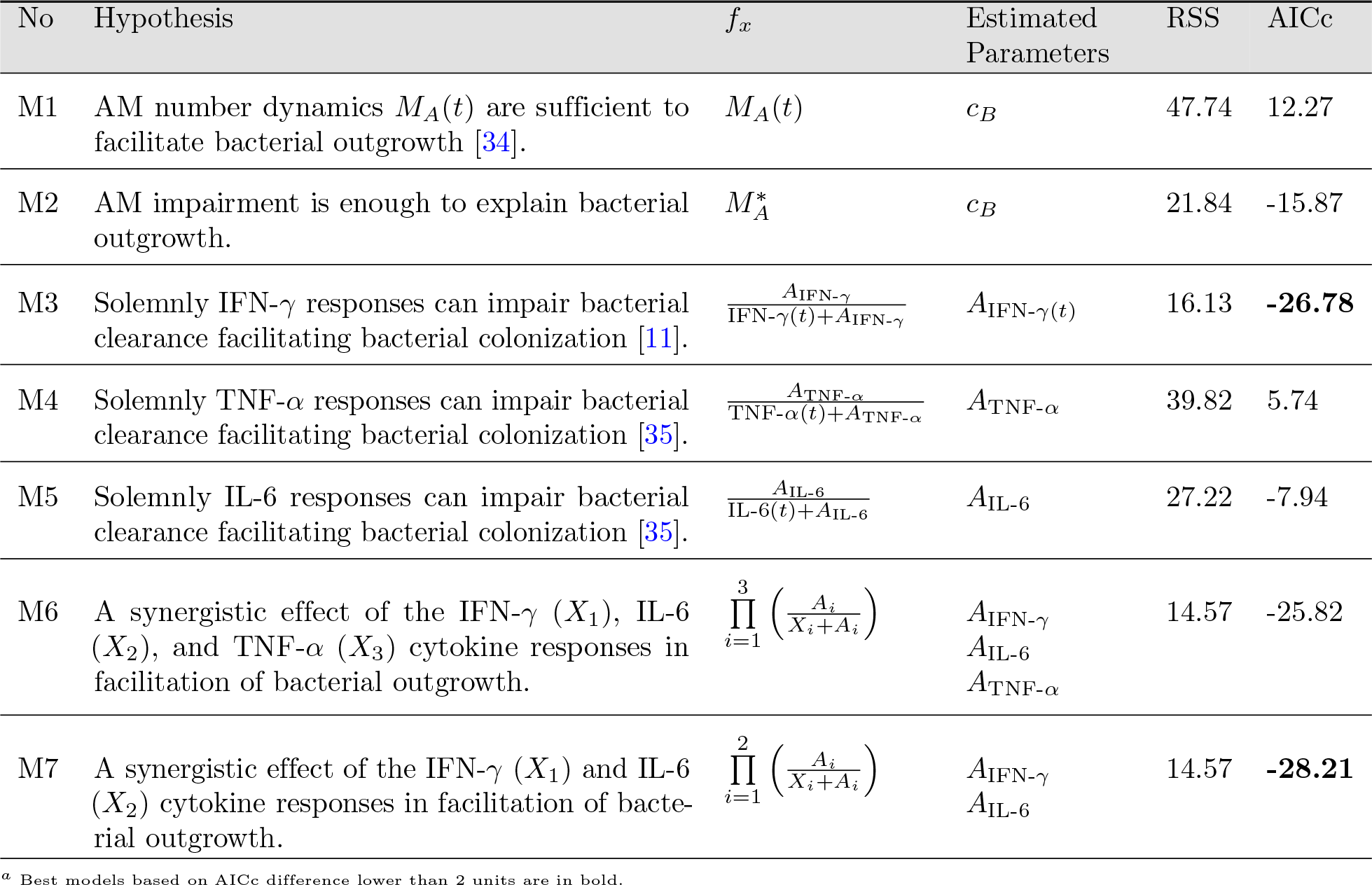
Selected list of the coinfection models to test different hypotheses that facilitate *S. pneumoniae* colonization. A complete model list with estimated parameters can be found in Supplementary E.

**The IL-6 response contributes to the impairment of bacterial clearance in a temporal manner.** Both the models M6 and M7 suggested that IL-6 contributed to impairing bacterial clearance during coinfection. However, the AICc score for the model M5 pointed out that it may not be as crucial as IFN-*γ* for promoting bacterial growth. However, it could be deduced that IL-6 is involved in the majority of the best models (*e.g* M6, M7, M8 and M9), indicating a time-dependent detrimental role in curtailing bacterial outgrowth.

To disentangle the time-dependent roles of significant pro-inflammatory cytokines, we selected the model M6 to perform *in silico* experiments. Simulation results suggested that a single neutralization of IFN-*γ* directly modulates the bacterial clearance. In contrast, neutralization of IL-6 or TNF-*α* did not present a conclusive role in impairing bacterial clearance (Figure 5a). Interestingly, when the IFN-*γ* response was neutralized in the model, simulations suggested that the IL-6 response may increase the duration of bacterial colonization in the lungs (Figure 5a). This possibly explains the experimental observation that coinfected mice still presented a marginal increase in the bacterial burden and degree of lethality even after *in vivo* IFN-*γ* blockade [11].

**Figure 5.**
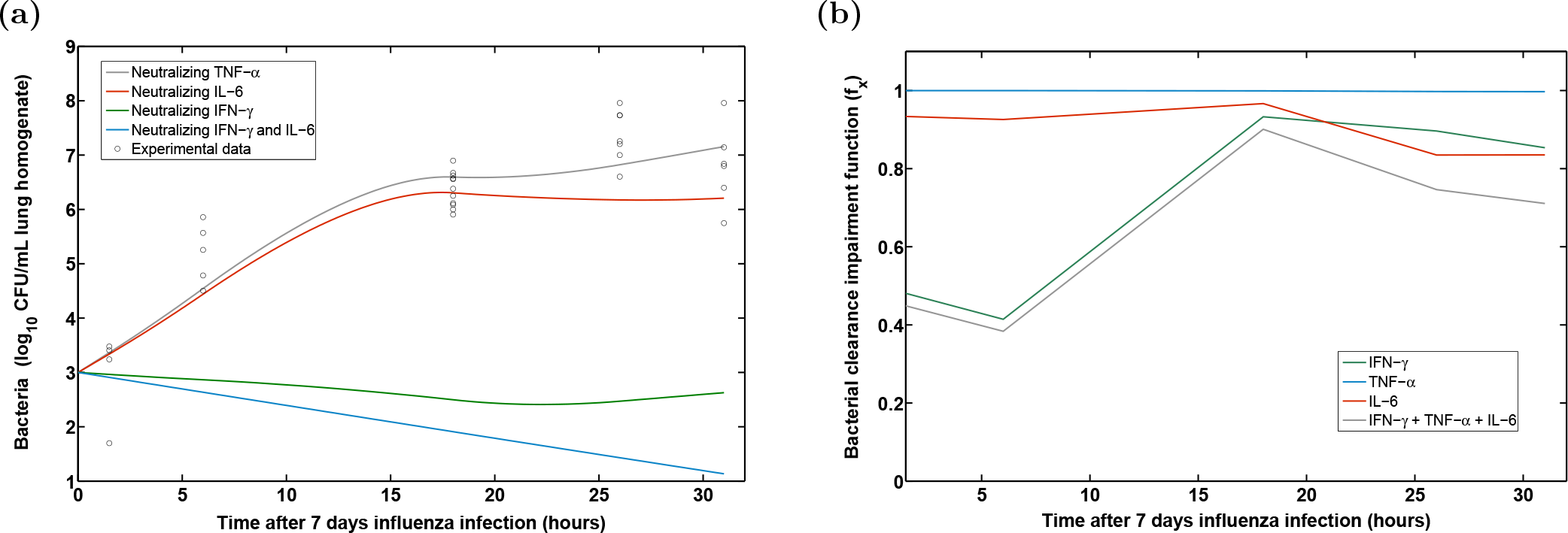
Simulations for the coinfection model M6. *In silico* neutralizations of the different pro-inflammatory responses in model M6 are presented in the panel (a). The time-dependent contributions of pro-inflammatory cytokines to the impairment of bacterial clearance by the function *f_x_* are depicted in the panel (b). When the mathematical function *f_x_* is 1 means that there is no any impairment to the bacterial clearance.

To weigh the inhibitory effects of different pro-inflammatory cytokines on bacterial clearance, we simulated the time evolution of the bacterial clearance impairment function (*f_x_*) for the different pro-inflammatory cytokines using the model M6. Indeed, the main modulator early after *S. pneumoniae* coinfection was IFN-*γ*. However after 18 hours the detrimental effects of IL-6 for the impairment of the bacterial clearance became more apparent (Figure 5b). In contrast, model selection procedures and numerical simulations rejected the hypothesis that TNF-α kinetics contributed to the impairment of bacterial clearance yielding one of the worst models (Table 1). These *in silico* results were supported by other model structures (M7, M8 and M9 at Supplementary E) and parameter uncertainty studies (Supplementary F).

**Stochastic fate decision explains the dichotomy in bacterial burdens observed in *S. pneumoniae* infection but plays only a minimal role during coinfections.** The bacterial load detected in the BAL and lung tissue revealed that the single *S. pneumoniae* infected animals divided into two groups at 18 hpi (Figure A.1). We found that 50% of the animals had cleared the bacteria while the remaining 50% were still colonized at this time point. This dichotomy was also reported by Smith *et al.* [30] and was suggested to result from the heterogeneity of the biological host.

**Figure 6.**
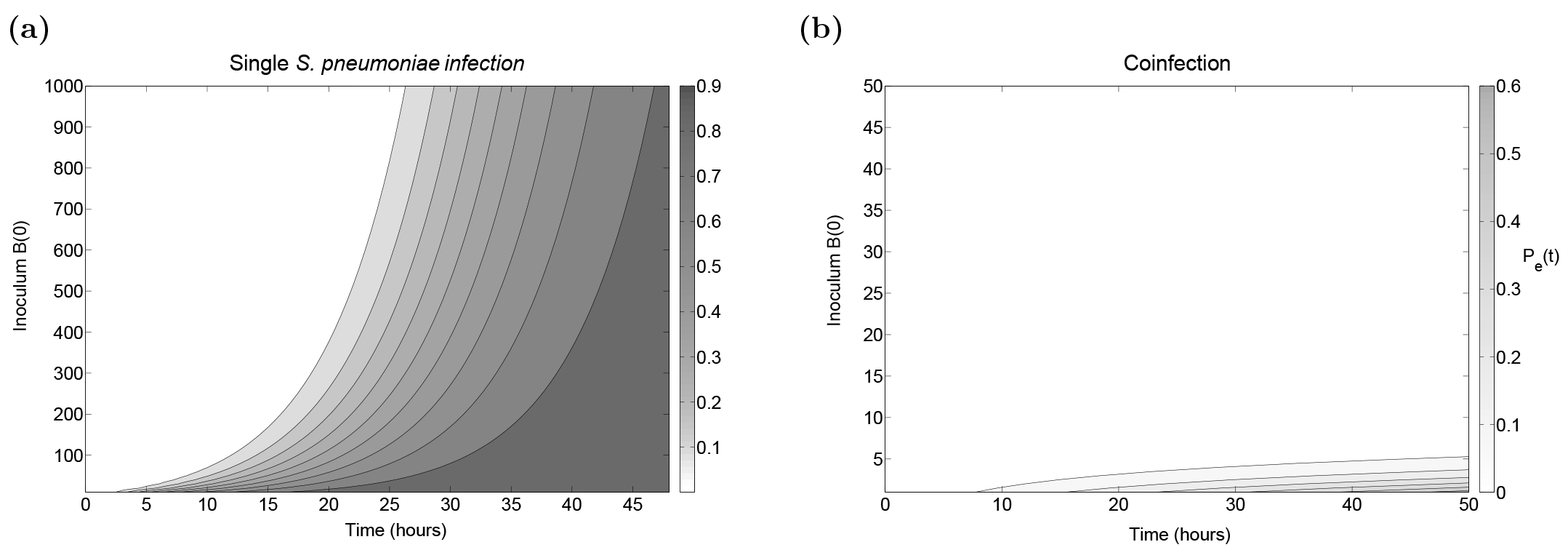
Probability of *S. pneumoniae* extinction dynamics. The branching process formulation governed by the Kolmogorov equation (Supplementary C) provides the probability of eradication *P_e_*(*t*) at time t for single *S. pneumoniae* infection and coinfection considering parameter values from model D2 (Supplementary D) and model M2 respectively.

To evaluate the stochastic fate decision, we derived the probability of extinction by developing a stochastic Markov branching process analogous to the model (1), see Supplementary C. For the single *S. pneumoniae* infection, the probability of bacterial extinction at 35 hpi is approximately 50% (Figure 6a), supporting the hypothesis of cell to cell heterogeneity [30]. In contrast, a stochastic fate decision with cell to cell heterogeneity is suggested to play only a minimal role in the case of coinfection (Figure 6b).

**Impairment of bacterial clearance by direct IAV kinetics is not supported.** Viral titers in the single IAV infected group remained stable at day 7 post the viral infection, *i.e* 1.5 and 6 hpi post secondary challenge, and the viral load then declined from 26 hpi (see Supplementary Figure A.4b). In accordance with previous experimentalobservations by Smith *et al.* [30], viral titers of the coinfected group showed a marginal, however not statistically significant viral rebound at 31 hpi, as the coinfected mice yielded higher viral loads than the single infected mice.

The modelling work by Smith *et al.* [30] tested the hypothesis that influenza infection modulates the bacterial clearance by AM phagocytosis. However, their model did not include aspects of host defense [30]. As a comparison, we considered the inhibitory function proposed by [30] using the kinetics of the viral load detected in our experiments (model M10 of Supplementary E). Model selection procedures however did not support the hypothesis that a time-dependent modulation by IAV kinetics contributes to the impairment of bacterial clearance.

**The modulation of nutrient availability through the IAV infection promotes the extent of initial bacterial colonization but not the decision of replication.** Experimental evidence by Siegel *et al.* [8] established that higher rates of disease during coinfection could stem from increased sialic acid availability that further supports bacterial colonization and proliferation. From a mathematical point of view, nutrient sources can be represented by the current capacity *K_B_* in equation (1). By the steady state and stability analysis presented in the Supplementary B, which is independent of parameter fitting procedures, it can be inferred that the bacterial nutrient source (*K_B_*) determines the size of the initial bacterial colony but not the decision to grow. Of note, this approach is also valid assuming more complex logistic growth terms. For instance, the mathematical term 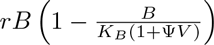 previously proposed by Smith *et al.* [30] to integrate the increase in the carrying capacity, *K_B_* (1 + Ψ*V*) may provide equivalent conclusions.

## Discussion

The threat of newly emerging pandemic IAV strains together with the increasing prevalence of antibiotic-resistant bacterial pathogens underline the need for a complete understanding of the mechanisms for secondary pneumonia. Recently, a substantial number of studies have uncovered several mechanisms through which IAV compromises efficient anti-bacterial defense. These different contributing factors for the viral-bacterial synergism highlight the multifactorial nature of copathogenesis.

At times however, the identified mechanisms are contradictory, *e.g* both de-sensitization and hyper-sensitization have been reported to favor bacterial infection post-influenza infection [9, 12, 36]. Nevertheless, a common observation is an altered pro-inflammatory cytokine response to the bacterial infection if preceded by influenza and many studies have attributed the altered lung cytokine milieu to be sufficient to skew host susceptibility to severe secondary pneumococcal infection [10, 11, 37].

The relative contribution of the different identified players however has not been addressed comprehensively to date. Although murine experiments are crucial to advance our knowledge of mechanisms which underlie virus-bacteria interactions, examining all different factors in detail renders an extremely difficult work. In this study, we combined an *in vivo* experimental approach and mathematical modelling to dissect the contributions of some of the circulating hypotheses proposed in driving bacterial outgrowth in IAV pre-infected hosts with the main focus on pro-inflammatory cytokines.

Unlike described by previous reports, in our experiments the inflammatory responses to secondary pneumococcal infection were not dampened but instead heightened when compared to the single bacterial infection [9]. Through mathematical modelling we identified a definite role for IFN-*γ* in impairing anti-pneumococcal clearance leading to outgrowth and systemic spread. This was in line with the finding that IFN-*γ* released during IAV-infection suppresses alveolar macrophage phagocytosis and increases oxidative radicals by downregulating their expression of the scavenger receptor MARCO in a state of coinfection, favoring bacterial outgrowth [9, 11, 36]. As *in vivo* neutralization of IFN-*γ* failed to fully alleviate bacterial outgrowth following coinfection with IAV and *S. pneumoniae*, other mediators are most likely to also contribute to the strongly impaired bacterial clearance [11].

In accordance with other possible key players during coinfection, we found in our murine experiments significantly higher concentrations of IL-6 and TNF-*α* during coinfection compared to the single infections [12, 37]. The exact role of these cytokines in the pathogenesis of coinfections still remains debatable. Here, our *in silico* studies suggest that the inhibitory effect of IL-6 is both concentration and time-dependent but not as conclusive for bacterial outgrowth as IFN-*γ*. Early after pneumococcal infection, IL-6 is likely to play a predominantly protective role due to its immune-regulatory function in the feedback circuit of cytokines. Indeed, it was reported that IL-6 KO mice displayed a significantly enhanced susceptibility to *S. pneumoniae* infection with elevated concentrations of TNF-*α*, Interleukin 1,β (IL-1β) and IFN-*γ* in the lungs in comparison to wild-type mice [38]. On the other hand, due to its potent pro-inflammatory functions, IL-6 is also known to be a marker for disease severity in pneumococcal infections [39]. Importantly, our finding that increased amounts of IL-6 predominantly impair bacterial clearance in synergism with the IAV-dependent IFN-*γ* present in the coinfected lung reflects how alterations in the inflammatory status affect host susceptibility in a dynamic temporal manner.

In order to address the possibly overlapping roles of IFN-*γ* and IL-6, we observed that TNF-*α* showed the least contribution in impairing bacterial clearance following influenza. This was well in line with previous findings describing that TNF-*α* neutralization elevated mortality during single *S. pneumoniae* infection and IAV coinfections suggesting more of a protective role for TNF-*α* [40,41]. However, TNF-*α* is in contrast known to induce apoptosis of various pulmonary cells and disrupt the epithelial barrier integrity and therefore TNF-*α* blockade ameliorates pulmonary immunopathology in single IAV infected animals [42–44]. In our mathematical models however, effects such as the detrimental contribution of immunopathology on *S. pneumoniae* outgrowth were not considered. Therefore it is likely that an excessive production of TNF-*α* during coinfections may potentiate host tissue damage and thereby may still exert an indirect negative influence on bacterial clearance.

Under homeostatic conditions, tightly regulated immune responses are co-ordinated by the action of pro-and anti-inflammatory cytokines and chemokines with the goal to clear pathogens and at the same time curtail immunopathology. By our experimental and mathematical modelling approach we found that during coinfections this balance is lost due to the exaggerated amounts of pro-inflammatory cytokines causing a pathogenic effect on bacterial clearance. Notably, this finding was further supported by the results of challenging our mathematical models for the IAV-S. *pneumoniae* coinfection with the cytokine data from the single *S. pneumoniae* infection group. Here, *in silico* results predicted bacterial clearance as in the single bacterial infection without further parameter fitting (Figure E.1).

In conclusion, by combining tailored experimental data and mathematical modelling our study clearly demonstrated a strong detrimental effect of IFN-*γ* alone and in synergism with IL-6 but no conclusive pathogenic effect of IL-6 and TNF-*α* alone. *In silico* knock-out predictions pave the way for further murine experiments to proof the advantage of a double IFN-*γ* and IL-6 neutralization approach. Ultimately, these findings correlate well with our previous results [29] suggesting that the increased levels of pro-inflammatory cytokines (the “inflammaging” state), in particular IFN-*γ*, contribute to the reported impaired responses in people over 65 years of age. Thereby, IFN-*γ* plays a pivotal role in driving severe disease during primary IAV infection in the elderly as well as bacterial outgrowth during coinfections.

In summary, our experimental setting and subsequent modelling approach suggest hierarchical and dynamic effects of several hallmark innate immunological factors on pneumococcal outgrowth in IAV pre-infected hosts. On this basis, future experimental studies will need to dissect the exact mechanisms underlying these detrimental effects predicted by our models. Our study also underscores the importance of further determining immune cell function, especially for AMs. Ultimately, such full understanding of the downstream effects of the altered inflammatory response in bacterial coinfection following IAV will be crucial when attempting to design future prophylactic and therapeutic interventions.

## Materials and Methods

### Mice

7-8 weeks old wildtype C57BL\6J female mice were purchased from Harlan Winkelmann (Borchen, Germany) and housed in specific pathogen-free conditions. All animal experiments were approved by the local ethical body “Niedersächsisches Landesamt für Verbraucherschutz und Lebensmittelsicherheit”.

### Viral and bacterial pathogens

For viral infection experiments, Influenza A virus IAV strain A/Puerto Rico/8/1934 (A/PR8/34) subtype H1N1 was used [31]. The virus was grown and stored as described previously in [31]. The tissue culture infectious dose (TCID_50_) was calculated using the Reed and Muench endpoint calculations. For bacterial infection experiments *S. pneumoniae* serotype 4, TIGR4 (ATCC BAA-334^TM^) was used. Bacteria were grown to mid-logarithmic phase in Todd-Hewitt yeast medium (THY; THB Sigma-Aldrich, Germany and Yeast extract, Roth, Germany) at 37°C for the preparation of frozen stocks. Bacterial counts were determined by plating 10-fold serial dilutions on blood agar plates (BD Diagnostic Systems, Columbia Agar with 5% sheep blood) overnight for 16-18 h at 37°C with 5% CO_2_. Before each infection, aliquots were thawed, centrifuged at 8000 rpm and resuspended in the desired amount of phosphate buffered saline (PBS, Gibco, UK) followed by serial plating to confirm the infection dose.

### Infection experiments

Mice were sedated via intraperitoneal injection of ketamine/rompun solution (working concentration of 0.1ml/10g per mouse). For viral challenges, a sub-lethal dose of 31 TCID_50_ was administered intranasally in 25*μ*l PBS. For bacterial challenges, mice were held with their backs on an intubation slope, and an infectious dose of 1 × 10^6^ CFU in 25*μ*l PBS was instilled at the end of their nasopharynx using a long flexible pipet tip. All the mice were monitored and scored for the following clinical symptoms: weight loss, mobility, posture, pilo-erection, respiration, response to stimuli and eye infection. An animal with a severe score of any one of the parameters or moderate scores for two-three parameters was euthanized.

### Determination of bacterial colony forming units in blood, bronchoalveolar lavage and lung tissue

Bacterial colony forming units (CFU) were determined post-mortem by plating 10-fold serial dilutions of the blood, BAL and lung tissue homogenate. For the blood, 5 *μ*L of heart blood were collected and diluted in 45*μ*l PBS. BAL samples were obtained by flushing the lung once with 1 mL PBS. Next, the lungs were perfused with PBS and excised. Whole lungs were homogenized in 1 mL PBS though a 100 *μ*m cell stainer (Corning Inc, USA). Serial dilutions of BAL and lung homogenates were prepared and 10*μ*l were plated onto blood agar plates. Plates were incubated overnight at 37°C and 5% CO_2_. For determination of CFU the colonies of each dilution were counted manually and calculated as CFU/mL.

### Quantification of viral load with qRT-PCR

Lungs were perfused with PBS and the RNA was extracted from lung tissue homogenates with the RNeasy mini kit (Qiagen, Germany) according the manufacturer’s protocol. After extraction and purification, cDNA synthesis was performed with 100 ng RNA using the first strand synthesis buffer (Invitrogen, USA). For cDNA quality control, a PCR of the housekeeping gene RPS9 (Eurofins MWG, Germany) was performed. Viral nucleoprotein copy numbers were quantified by RT-PCR on a Light cycler (Roche) using a plasmid standard with defined NP copy numbers. The following primers were used: rps9 5’CTGGACGAGGGCAAGATGAAGC and 3’TGACGTTGGCGGATGAGCACA; np 5’GAGGGGTGAGAATGGACGAAAAAC and 3’CAGGCAGGCAGGCAGGACTT.

### Flow cytometry for alveolar macrophages

Single cell suspensions of perfused excised lungs were prepared by enzymatic digestion for 45 minutes at 37°C with Iscove’s Modified Dulbecco’s Medium (IMDM) containing GlutaMax-1 (Gibco, USA) supplemented with 5% fetal bovine serum (FBS; PanBiotech, Germany), 0.2 mg/ml collagenase D (Roche, Germany) and 1 mg/ml DNAse (Sigma-Aldrich, Germany). The enzymatic reaction was stopped with 5mM EDTA and cell suspensions were filtered through a 100 *μ*m strainer.

Cells were pelleted by centrifugation and resuspended in LIVE/DEAD^®^ fixable blue stain (ThermoFisher, USA) and anti-mouse CD16/CD32 antibody (purified; BioLegend, USA) as Fc-block following erythrocyte lysis. Cell surface marker staining for lung resident alveolar macrophages (SSChighFSChighCD11b-F4/80+autoflourescence+ AM) was performed using anti-mouse CD11b (Pacific Blue; BioLegend, USA) and anti-mouse F4/80 (PE-Cy7; BioLegend, USA). All samples were acquired on a BD LSRII Fortessa instrument with FACS DIVA software (BD) and analyzed using FlowJo software (Tree Star).

### Cytokine measurement

The protein concentration of the cytokines IFN-*γ*, TNF-*α*, IL-6, IFN-*β*;, IL-22 and the chemokines MCP-1 and GM-CSF was measured with a customized Mouse LegendPlex*^™^* kit (BioLegend, USA) according to the manufacturers protocol. All samples were acquired on a BD LSRII Fortessa with FACS DIVA software (BD) and analyzed with the provided LegendPlex*^™^* Data Analysis Software (BioLegend, USA).

### Mathematical Models

This work advocates to disentangle the mechanisms modulated by IAV that can contribute to the colonization of *S. pneumoniae*. Although, we recognize that bacterial infection may enhance viral release from infected cells [30], these interactions are outside of the aim of the work proposed here. Therefore, we model the *S. pneumoniae* infection (B) with the following ordinary differential equations (ODEs):

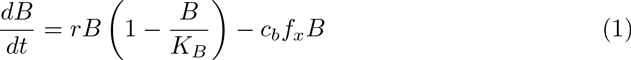

where *r* is the bacterial proliferation rate with a maximum carrying capacity *K_B_*. Phagocytosis of the bacteria is considered by the multiplicative term c_b_*f_x_*, where c_b_ is the constant phagocytosis rate and the term *f_x_* is the mathematical function which served to test different hypotheses.

The previous work by Smith *et al.* [30] proposed and tested that IAV induces phenotypic changes in AM. However, the work in [30] assumed the restrictive condition that thephagocytosis rate *c_b_* is impaired directly by the viral load (*V*(*t*)). This hypothesis was written with the term *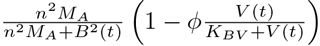* where the values for the uptake of bacteria by AM were expressed by *n* and the number of AM was constant in the quasi steady state 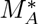[30]. The reduction of the bacterial clearance was included by the saturation function *ϕV*(*t*)/(*K_BV_+V*(*t*)), where *ϕ* is the maximal reduction of the phagocytosis rate and *K_BV_* is the half-saturation constant.

In order to evaluate the AM dynamics, our model section considers AM experimental data to build a piecewise linear function to feed the model term *M_A_*(*t*). Additionally, we tested the hypothesis that AM depletion by IAV is a sufficient mechanism to facilitate bacterial outgrowth. This is represented by a direct input of 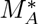 in equation (1).

Beyond the work of [30], different functions (*f_x_*) were tested to dissect the contributions of pro-inflammatory cytokines (IFN-*γ*, TNF-*α*, and IL-6) to modulate the bacterial clearance. To this end, we used the experimental data for the different cytokines to build piecewise linear functions to dynamically feed the equation (1). We opted for this approach instead of mechanistic modelling with ODEs for each cytokine for different reasons. First, our main aim was not to quantify dynamics but to determine the contribution of important pro-inflammatory cytokines in promoting bacterial colonization. Second, modelling with ODEs for each cytokine may not improve the model selection process but only increase the complexity of mathematical models and parameter fitting procedures. Finally, our experimental data sets were measured frequently enough allowing us to use the data as an input in the equation (1).

To evaluate if IAV infection modulates a specific pro-inflammatory cytokine *X*(*t*), which subsequently facilitates bacterial colonization, we considered the saturating term 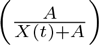 with *A* as the “half-saturation constant”. Different mathematical terms and other term combinations to dissect the pro-inflammatory effect responsible to impair bacterial clearance are summarized in Table 1 and Supplementary E.

### Parameter Fitting

Note that the purpose of our *in silico* work was not biological quantification of the experiments but using a model selection process to provide mechanistic insights into coinfection. Thus, in this case, finding a working set of parameter values deemed sufficient. Nevertheless, for good practice procedures, we checked identifiability and parameter uncertainty expanded in the Supplementary F.

To focus on the mechanisms that promote *S. pneumoniae* colonization, we fixed the growth rate (*r* =1.13 h^-1^) and the carrying capacity (*K_B_* =2.3×10^8^ CFU/ml) corresponding to single *S. pneumoniae* infection from previous works [18]. Using similar reasoning as Smith *et al.* [18], we assumed that only a proportion of the bacterial inoculum may reach the lung since some bacteria can be removed by mucocilliary mechanisms. Thus, we considered 1000 CFU/mL *S. pneumoniae* as inoculum to fit the model parameters in (1). Note that different assumptions of parameters and inoculation will only rescale the fitted parameters, but not the mechanistic insights from the model selection procedures. Other model parameters were obtained minimizing the residual sum of squares (RSS) between the model output and the experimental measurement, both on log scales. ODEs were solved in Matlab software using the ode45 solver. The minimization of RSS was performed using different optimization solvers, including both deterministic and stochastic methods. However, the Differential Evolution algorithm (DE), a global optimization algorithm, was selected to avoid relying on any initial parameter guesses and producing more robust results than other tested methods [45–47]. In separate form, model fitting was performed to all the models summarized in Table 1 for the single infection (T4) and the IAV+T4 coinfection starting at day 7 post IAV infection respectively.

### Model Selection

The Akaike information criterion (AIC) was used to compare the goodness-of-fit for models that evaluate different hypotheses [48]. A lower AIC value means that a given model describes the data better than other models with higher values. However, small differences in AIC scores (*e.g. <2*) are not significant [48]. For a small number of data points, the corrected (AICc) has the form:

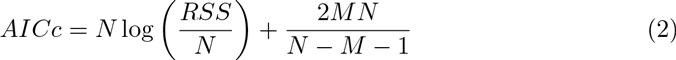

where *N* is the number of data points, *M* is the number of unknown parameters and *RSS* is the residual sum of squares obtained from the fitting routine.

### Parameter uncertainty

Bootstrapping is a statistical method for assigning measures of accuracy to estimates [49, 50]. Due to the large variability of viral and bacterial infections, bootstrapping methods have been applied to different viral infectious diseases [47, 51]. Here, the non-parametric bootstrap was considered. Of note, instead of using all replications at each time point, three replications at each time point were randomly selected and used to generate a new data set. Note that this procedure was based exclusively on the observed measurement data. For each repetition, the model parameters were refitted to obtain the corresponding parameter distribution. The 95% Confidence Interval (CI) of parameter estimates was computed using the outcome of the bootstrap method [50]. For each parameter, the 2.5% and 97.5% quantiles of the estimates were used to form the 95% CI.

### Parameter identifiability

A relevant aspect to verify in quantifying mechanisms is whether model parameters are identifiable based on the measurements of output variables [52–55]. A system that is algebraically identifiable may still be practically non-identifiable if the amount and quality of the measurements is insufficient and the data shows large deviations. The computational approach proposed in [56] exploits the profile likelihood to determine identifiability and was considered here. This method is able to detect both structurally and practically non-identifiable parameters. Briefly, the idea behind this approach is to explore the parameter space for each parameter *θ_i_* by re-optimizing the RSS with respect to all other parameters *θ_j≠i_*. The main task is to detect directions where the likelihood flattens out [56]. The resulting profiles are plotted versus each parameter range with the respective 95% CI to assess the parameter identifiability.

### Stochastic Modelling

In order to evaluate the stochastic fate between different cells in single *S. pneumoniae* infection and coinfection, we considered the stochastic Markov branching process. This framework has been applied to predict immune response dynamics, for instance for CD8+ T cell clonal expansion [57, 58] and infectious diseases [59, 60]. According to the Markov branching process definition, each *S. pneumoniae* cell can proliferate at rate *r* and die at rate *c_b_* (birth and death process). We derived an analytical solution for the extinction probability information to investigate the extinction probability evolution for different inocula *B*(0). Further details of the stochastic modelling framework can be found in Supplementary C.

### Statistical Analysis

All statistical analyses were performed by a non-parametric t-test called Mann-Whitney using the Graph Pad Prism software (Graph Pad Software, La Jolla/USA).

## Acknowledgments

This work was supported by iMed-the Helmholtz Initiative on Personalized Medicine and by the Department of Systems Immunology at the Helmholtz Centre for Infection Research. Dunja Bruder was supported by the President’s Initiative and Networking Fund of the Helmholtz Association of German Research Centers (HGF) under contract number W2/W3-029 and by a grant from the German Research Foundation (DFG) in frame of the SFB854 (subproject A23N). NS received support from the International Research Training Group (IRTG) 1273.

